# Identification of M-Sec as a unique cellular regulator of CSF-1 receptor activation

**DOI:** 10.1101/2024.09.03.610939

**Authors:** Randa A. Abdelnaser, Masateru Hiyoshi, Naofumi Takahashi, Youssef M. Eltalkhawy, Hidenobu Mizuno, Shunsuke Kimura, Koji Hase, Hiroshi Ohno, Kazuaki Monde, Akira Ono, Shinya Suzu

**Affiliations:** Joint Research Center for Human Retrovirus Infection, Kumamoto University, Kumamoto, Japan; Research Center for Biological Products in the Next Generation, National Institute of Infectious Diseases, Tokyo, Japan; International Research Center for Medical Sciences, Kumamoto University, Kumamoto, Japan; Division of Biochemistry, Faculty of Pharmacy, Keio University, Tokyo, Japan; Laboratory for Intestinal Ecosystem, RIKEN Center for Integrative Medical Sciences, Yokohama, Kanagawa, Japan; Department of Microbiology, Faculty of Life Sciences, Kumamoto University, Kumamoto, Japan; Department of Microbiology and Immunology, University of Michigan Medical School, Ann Arbor, MI, USA

## Abstract

Fms, the CSF-1 receptor encoding tyrosine kinase, is essential for tissue macrophage development, and the therapeutic target for many tumors. However, it is not completely understood how Fms activation is regulated. Here, we identify the cellular protein M-Sec as a unique regulator of Fms. In macrophages, Fms forms large aggregates via unknown mechanisms. We found that the inhibition or knockdown of reduced Fms aggregate formation and functional response of macrophages to CSF-1, which was consistent with reduced Fms activation after CSF-1 stimulation. When expressed in 293 cells, M-Sec augmented Fms aggregate formation and CSF-1-induced Fms activation. CSF-1 and M-Sec bind the cellular phosphatidylinositol 4,5-biphosphate (PIP2). The removal of PIP2-binding motif of Fms or M-Sec, or the depletion of cellular PIP2 reduced Fms aggregate formation. Moreover, M-Sec altered cellular distribution of PIP2. Since CSF-1-induced dimerization of Fms is critical for its activation, our findings suggest that M-Sec augments large Fms aggregate formation via PIP2, which brings Fms monomers close to each other and enables the efficient dimerization and activation of Fms in response to CSF-1.

## INTRODUCTION

Macrophages are innate immune cells that orchestrate homeostasis, inflammation or regeneration, and present across various tissues throughout the body (1, 2). The pool of macrophages in each tissue is maintained by the differentiation of bone marrow-derived monocytes and/or the self-renewal of macrophages originated from the precursors in extraembryonic yolk sac or monocytes in fetal liver (3). The development, proliferation and survival of tissue macrophages can be regulated by cytokines, such as CSF-1, IL-34 and CSF-2 (4). CSF-1 (also known as M-CSF) and IL-34 share the receptor Fms (5, 6), and mice lacking Fms are deficient in most tissue macrophages (7). Interestingly, mice lacking CSF-1 are deficient in most tissue macrophages, but not in Langerhans cells and microglia, whereas mice lacking IL-34 exhibit selective reductions in Langerhans cells and microglia, which is because CSF-1 is ubiquitously expressed whereas IL-34 is mainly expressed by keratinocytes and neurons (8, 9). Meanwhile, mice lacking CSF-2 (also known as GM-CSF) show functional defects in alveolar macrophages but no major deficiency in most tissue macrophages (10, 11). Thus, the CSF-1/IL-34-Fms axis is critical for the development of most tissue macrophages.

Macrophages are abundant immune cells in tumor microenvironment and the CSF-1/IL-34-Fms axis often induces pro-tumorigenic macrophages (12). Thus, Fms is one of the attractive therapeutic targets for a variety of tumors. For instance, the small molecule Fms inhibitor BLZ945 blocked the progression of glioma in preclinical models (13), and the anti-Fms antibody RG7155 reduced tumor burden in patients with diffuse-type giant cell tumors (14). Many clinical studies targeting Fms are ongoing (15).

Fms is a receptor tyrosine kinase and belongs to class III receptor tyrosine kinase that includes Flt3, Kit, PDGFRα, and PDGFRβ (16, 17). In the absence of CSF-1 or IL-34, Fms is present as a monomer and the Fms monomer is inactive due to cis-autoinhibition (16, 17). The binding of CSF-1 or IL-34, both of which are homodimers, induces the dimerization of Fms and releases the cis-autoinhibition (16–18). The change leads to the activation and auto-phosphorylation of Fms, which results in the tyrosine-phosphorylation of downstream proteins and activation of various signaling pathways, including MAP kinases (6, 17, 19, 20). Thus, the dimerization is critical for the subsequent activation and auto-phosphorylation of Fms. However, it is not completely understood how Fms dimerization is regulated. Interestingly, Fms monomers are clustered and form large aggregates in macrophages (21–23). The pre-formed Fms aggregates in which the monomers are close to each other may be beneficial for CSF-1 or IL-34 to dimerize and activate Fms (21). However, to what extent Fms aggregate formation contributes to dimerization/activation of Fms remains unexplored. Furthermore, little is known about molecular mechanism by which Fms forms the aggregates.

In this study, we show that a cellular protein M-Sec (also known as tnfaip2) is required for the Fms aggregate formation and important for an efficient activation of Fms in macrophages. M-Sec is highly expressed in macrophages and known to promote the formation of tunneling nanotubes (24, 25), the F-actin-containing long plasma membrane protrusions. Although we have demonstrated that M-Sec facilitates cell-to-cell transmission of human retroviruses, such as HIV-1 and HTLV-1 (26–28), physiological functions of M-Sec in macrophages other than the tunneling nanotube formation were not completely understood. Here, we provide evidence that M-Sec functions as a unique regulator of Fms activation.

## RESULTS

### M-Sec inhibition or knockdown reduces Fms aggregate formation in macrophages

As reported (21–23), Fms was detected as large aggregates in bone marrow-derived macrophages (Figure 1A, upper). In this study, we initially found that the large Fms aggregates were reduced by the M-Sec inhibitor (Figure 1A, lower), the small chemical that inhibits M-Sec-mediated tunneling nanotube formation (26, 29, 30). The size of Fms aggregates of M-Sec inhibitor-treated cells was smaller than that of the vehicle (DMSO)-treated control cells (Figure 1B), but their cell surface expression level of Fms was comparable (Figure 1C). The similar results were obtained when the macrophage cell line RAW264.7 was used: M-Sec knockdown reduced the formation of large Fms aggregates (Figure 1D) and the size of Fms aggregates (Figure 1E), but did not affect the cell surface Fms expression (Figure 1F).

**Figure 1.**
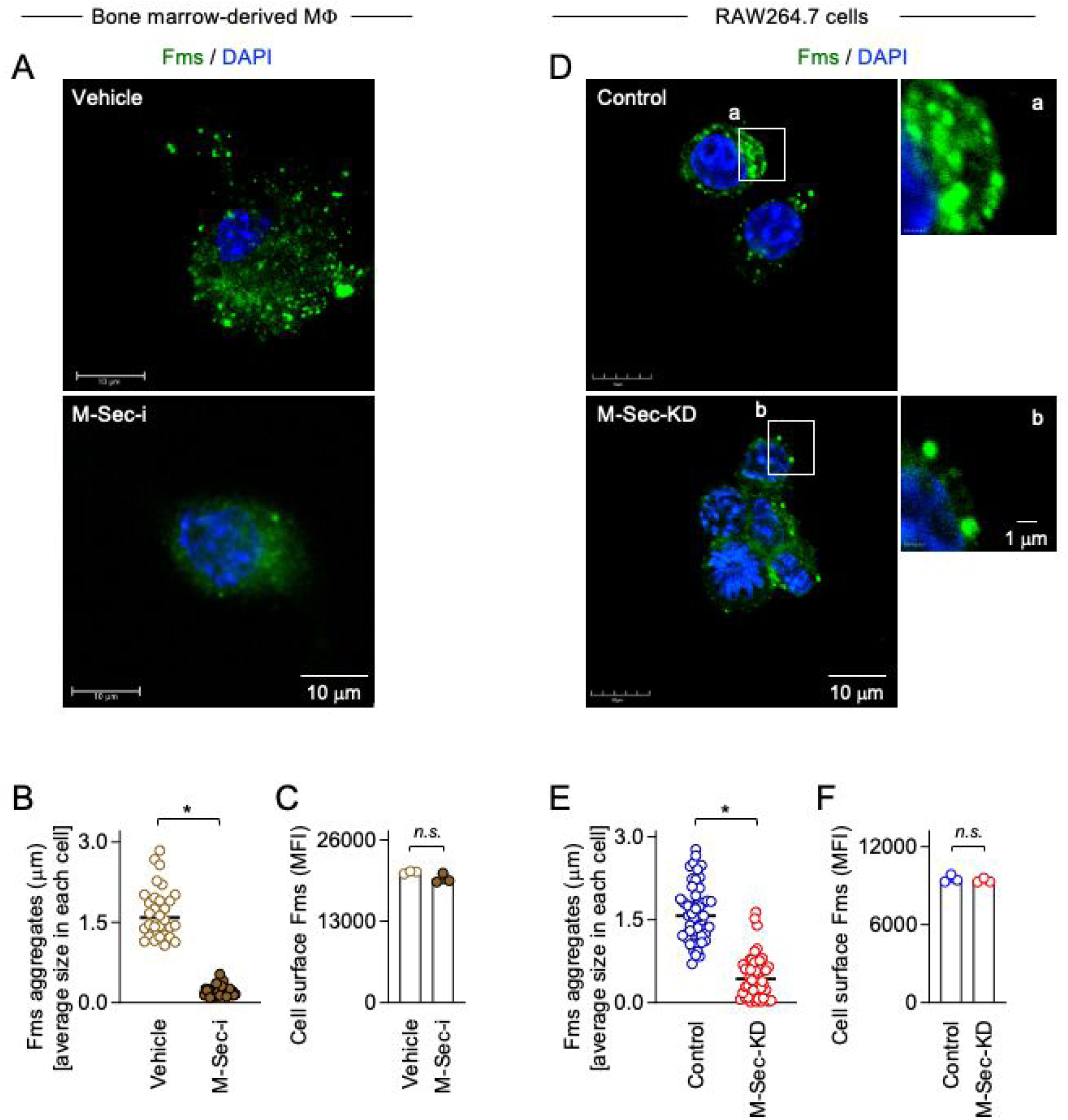
Effect of M-Sec inhibition or knockdown on Fms aggregate formation in macrophages. (A) Bone marrow-derived macrophages were treated with DMSO (Vehicle) or M-Sec inhibitor (M-Sec-i) for 48 hours, co-stained with anti-Fms antibody (green) and DAPI (blue), and analyzed by immunofluorescence. Scale bar: 10 μm. (B) Bone marrow-derived macrophages were analyzed as in **A**, and the average size of Fms aggregates in each cell is summarized (30 cells for each group). \**p* < 0.05. (C) Bone marrow-derived macrophages were treated with DMSO (Vehicle) or M-Sec inhibitor (M-Sec-i) for 48 hours, and analyzed for cell surface expression of Fms. The mean fluorescence intensity (MFI) is shown. *n.s.*, not significant. (D) The control or M-Sec knockdown (M-Sec-KD) RAW264.7 cells were co-stained with anti-Fms antibody (green) and DAPI (blue), and analyzed by immunofluorescence. Scale bar: 10 μm for left panels and 1 μm for right panels. (E) The control or M-Sec knockdown (M-Sec-KD) RAW264.7 cells were analyzed as in **D**, and the average size of Fms aggregates in each cell is summarized (50 cells for each group). \**p* < 0.05. (F) The control or M-Sec knockdown (M-Sec-KD) RAW264.7 cells were analyzed for cell surface expression of Fms. The MFI is shown. *n.s.*, not significant.

### M-Sec inhibition or knockdown reduces macrophage response to CSF-1, but not to CSF-2

We next examined how M-Sec inhibition or knockdown affects cellular response to CSF-1. When added to bone marrow-derived self-renewing macrophages (31), CSF-1 upregulated the expression of inducible nitric oxide synthase (iNOS) and CSF-2 upregulated more strongly (Figure 2A). Interestingly, the M-Sec inhibitor selectively reduced the iNOS upregulation by CSF-1 (Figure 2A). The similar result was observed for RAW264.7 cells: the M-Sec knockdown reduced the iNOS upregulation by CSF-1 (Figure 2B, left), but not that by CSF-2 (Figure 2B, right) or lipopolysaccharide (data not shown). Consistent with this, although CSF-1 and CSF-2 upregulated the production of nitric oxide, the M-Sec knockdown selectively reduced the nitric oxide upregulation by CSF-1 (Figure 2C).

**Figure 2.**
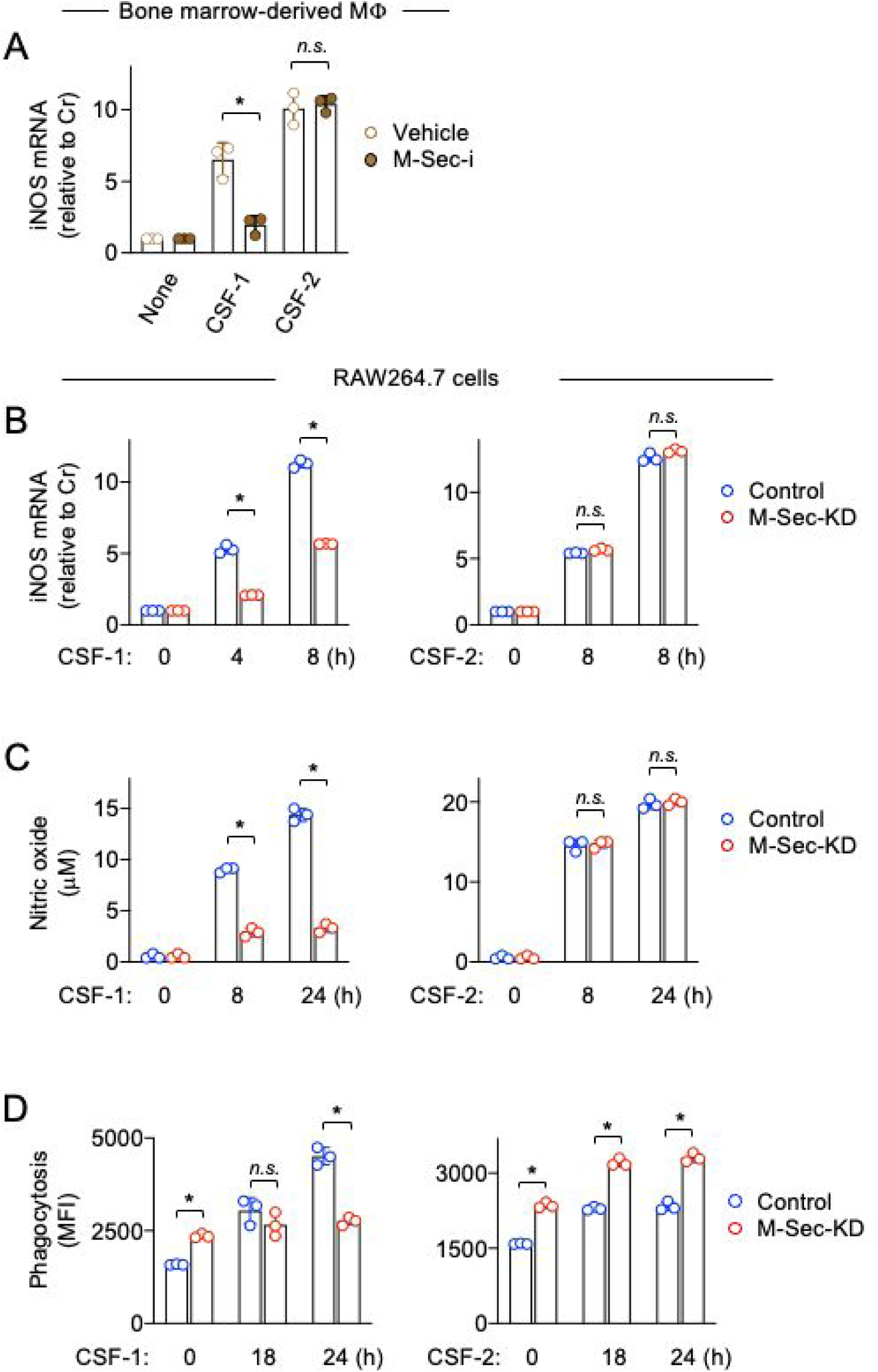
Effect of M-Sec inhibition or knockdown on functional response of macrophages to CSF-1. (A) Bone marrow-derived macrophages were treated with DMSO (Vehicle) or M-Sec inhibitor (M-Sec-i) for 24 hours, left untreated (None) or treated with CSF-1 or CSF-2 for 8 hours, and analyzed for iNOS mRNA expression by qRT-PCR (n=3). The expression level shown is relative to that of untreated cells. *n.s.*, not significant. \**p* < 0.05. (B) The control or M-Sec knockdown (M-Sec-KD) RAW264.7 cells were left untreated or treated with CSF-1 or CSF-2 for 4 or 8 hours, and analyzed for iNOS mRNA expression by qRT-PCR (n=3). The expression level shown is relative to that of untreated cells. *n.s.*, not significant. \**p* < 0.05. (C) The control or M-Sec knockdown (M-Sec-KD) RAW264.7 cells were left untreated or treated with CSF-1 or CSF-2 for 8 or 24 hours, and analyzed for the concentrations of nitric oxide in the supernatants using the Griess reagent (n=3). *n.s.*, not significant. \**p* < 0.05. (D) The control or M-Sec knockdown (M-Sec-KD) RAW264.7 cells were left untreated or treated with CSF-1 or CSF-2 for 18 or 24 hours, and analyzed for phagocytic activity by flow cytometry (n=3). The mean fluorescence intensity (MFI) is shown. *n.s.*, not significant. \**p* < 0.05.

The reduced response to CSF-1 by M-Sec knockdown was also observed in another assay: CSF-1 enhanced the phagocytic activity in the control cells, but not in the M-Sec knockdown cells, despite their higher basal phagocytic activity in the absence of CSF-1 (Figure 2D, left). Such differential response towards the control and M-Sec knockdown cells was not observed for CSF-2 (Figure 2D, right).

### M-Sec knockdown reduces MAP kinase activation in response to CSF-1, but not to CSF-2, in macrophages

We next examined how M-Sec knockdown affects signal activation in response to CSF-1. CSF-1 activated MAPKs (ERK, p38, and JNK) in the control cells, but weakly in the M-Sec knockdown cells (Figure 3A). Such differential response was not observed for CSF-2 (Figure 3B). The result was well consistent with the finding that the M-Sec knockdown reduced the cellular response to CSF-1, but not to CSF-2 (see Figure 2).

**Figure 3.**
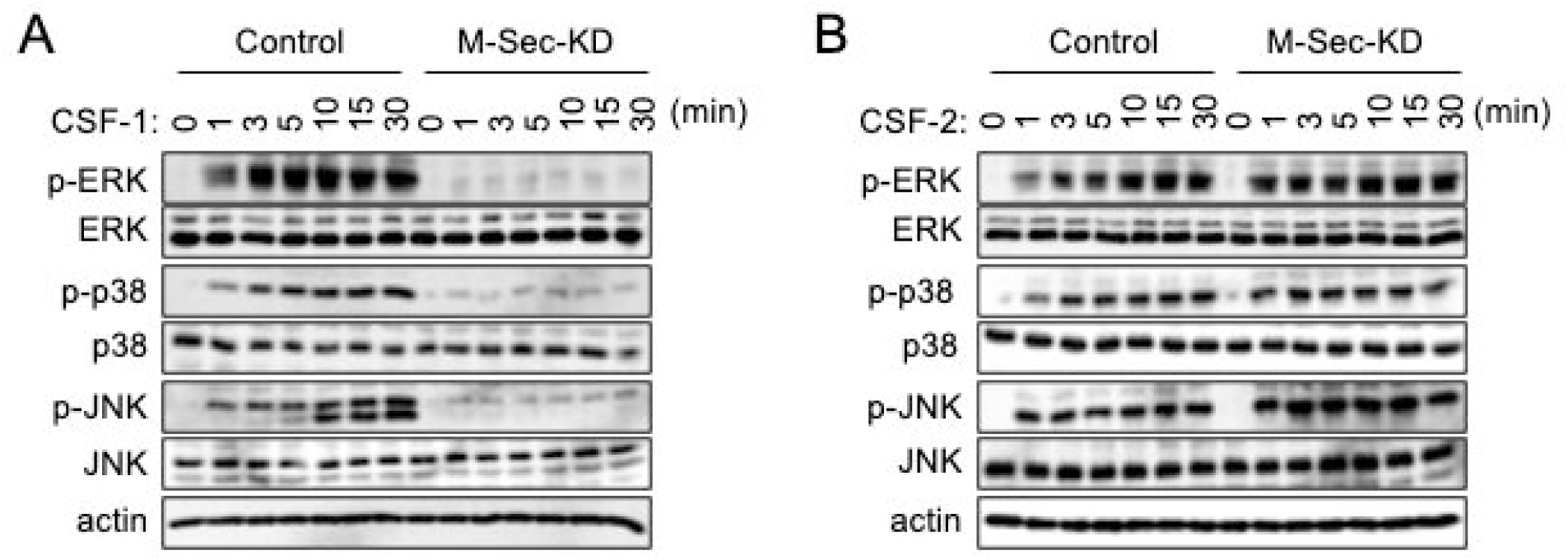
Effect of M-Sec knockdown on CSF-1-induced activation of MAP kinases in RAW264.7 cells. (**A**, **B**) The control or M-Sec knockdown (M-Sec-KD) RAW264.7 cells were serum starved for 12 hours, and left untreated or treated with CSF-1 (**A**) or CSF-2 (**B**) for the indicated periods. The total cell lysates were subjected to western blotting. Antibodies used were as follows: anti-phosphorylated ERK (p-ERK), anti-total ERK, anti-phosphorylated p38 (p-p38), anti-total p38, anti-phosphorylated JNK (p-JNK), and anti-total JNK. β-actin blot is the loading control.

The expression of human M-Sec in the M-Sec-knockdown mouse RAW264.7 cells cancelled their weak responses to CSF-1, including MAP kinase activation (Figure 4A), iNOS expression and nitric oxide production (Figure 4B), phagocytic activity (Figure 4C), and cell motility or osteoclastic differentiation (data not shown). Thus, the weak cellular response to CSF-1 by M-Sec knockdown was not due to off-target effects.

**Figure 4.**
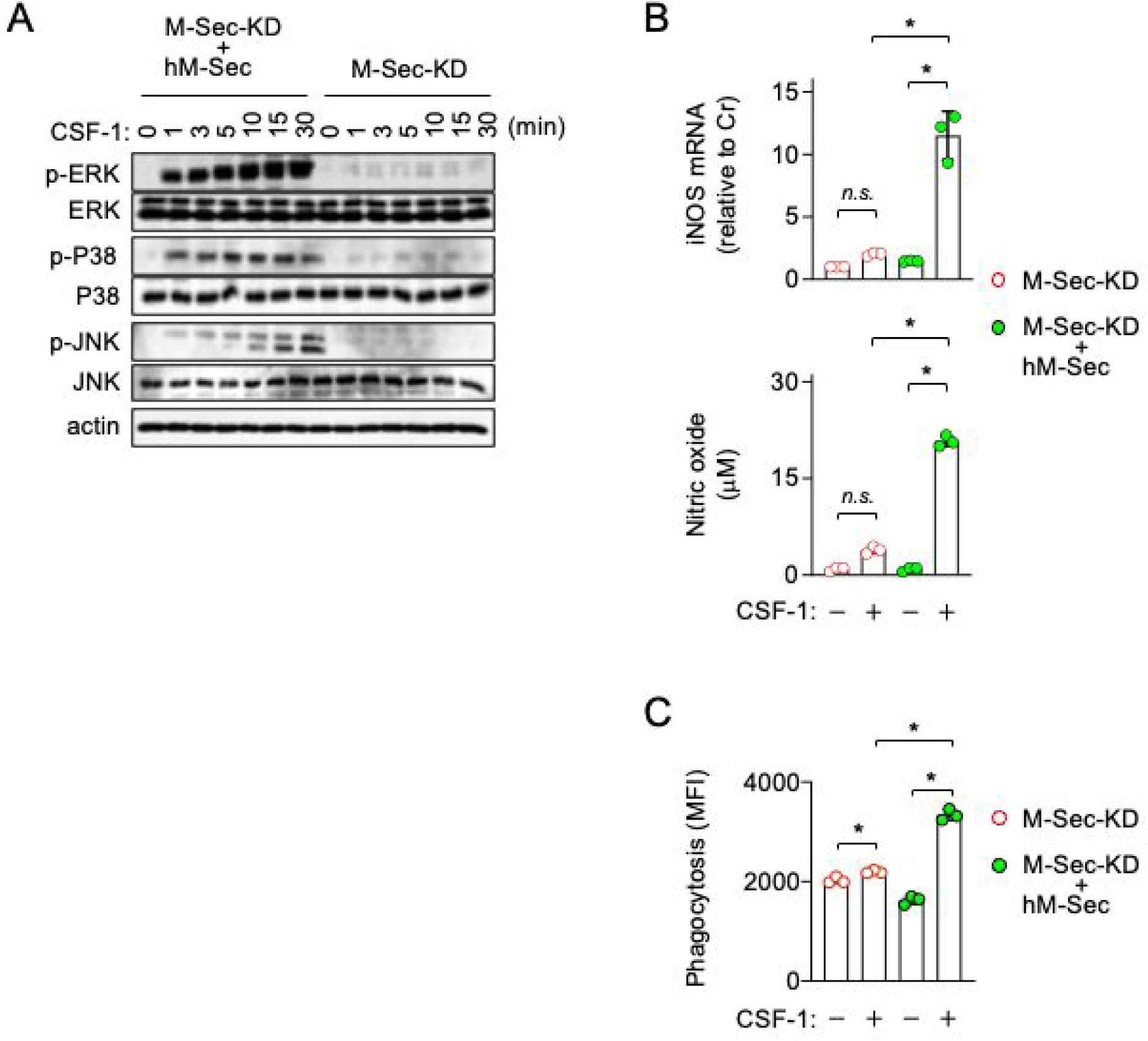
Effect of the exogenous expression of human M-Sec on the response of M-Sec knockdown RAW264.7 cells to CSF-1. (A) The M-Sec knockdown (M-Sec-KD) or human M-Sec-expressing M-Sec knockdown (M-Sec-KD + hM-Sec) RAW264.7 cells were serum starved for 12 hours, and left untreated or treated with CSF-1 for the indicated periods. The total cell lysates were subjected to western blotting. Antibodies used were as follows: anti-phosphorylated ERK (p-ERK), anti-total ERK, anti-phosphorylated p38 (p-p38), anti-total p38, anti-phosphorylated JNK (p-JNK), and anti-total JNK. β-actin blot is the loading control. (B) In the upper panel, the M-Sec knockdown (M-Sec-KD) or human M-Sec-expressing M-Sec knockdown (M-Sec-KD + hM-Sec) RAW264.7 cells were left untreated or treated with CSF-1 for 8 hours, and analyzed for iNOS mRNA expression by qRT-PCR (n=3). The expression level shown is relative to that of untreated M-Sec-KD cells. In the lower panel, the cells were left untreated or treated with CSF-1 for 24 hours, and analyzed for the concentrations of nitric oxide in the supernatants using the Griess reagent (n=3). *n.s.*, not significant. \**p* < 0.05. (C) The M-Sec knockdown (M-Sec-KD) or human M-Sec-expressing M-Sec knockdown (M-Sec-KD + hM-Sec) RAW264.7 cells were left untreated or treated with CSF-1 for 24 hours, and analyzed for phagocytic activity by flow cytometry (n=3). The mean fluorescence intensity (MFI) is shown. \**p* < 0.05.

### M-Sec inhibition or knockdown reduces CSF-1-induced Fms activation in macrophages

CSF-2 receptors consist of a unique α-chain and common β-chain (4), which are structurally unrelated to receptor tyrosine kinases including CSF-1 receptor Fms. Because the M-Sec inhibition or knockdown specifically reduced the cellular response to CSF-1, we next examined whether M-Sec inhibition or knockdown affects Fms activation. The activated Fms tyrosine-phosphorylates its downstream proteins (6, 17, 19, 20). In fact, CSF-1 induced the tyrosine-phosphorylated 150-300 kDa proteins in the vehicle-treated control bone marrow-derived macrophages, but such change was modest in the M-Sec inhibitor-treated cells (Figure 5A). The similar result was obtained when we compared the control and M-Sec knockdown RAW264.7 cells (Figure 5B, pTyr blot), which was consistent with the weak MAP kinase activation in response to CSF-1 in the M-Sec knockdown cells (see Figure 3A).

**Figure 5.**
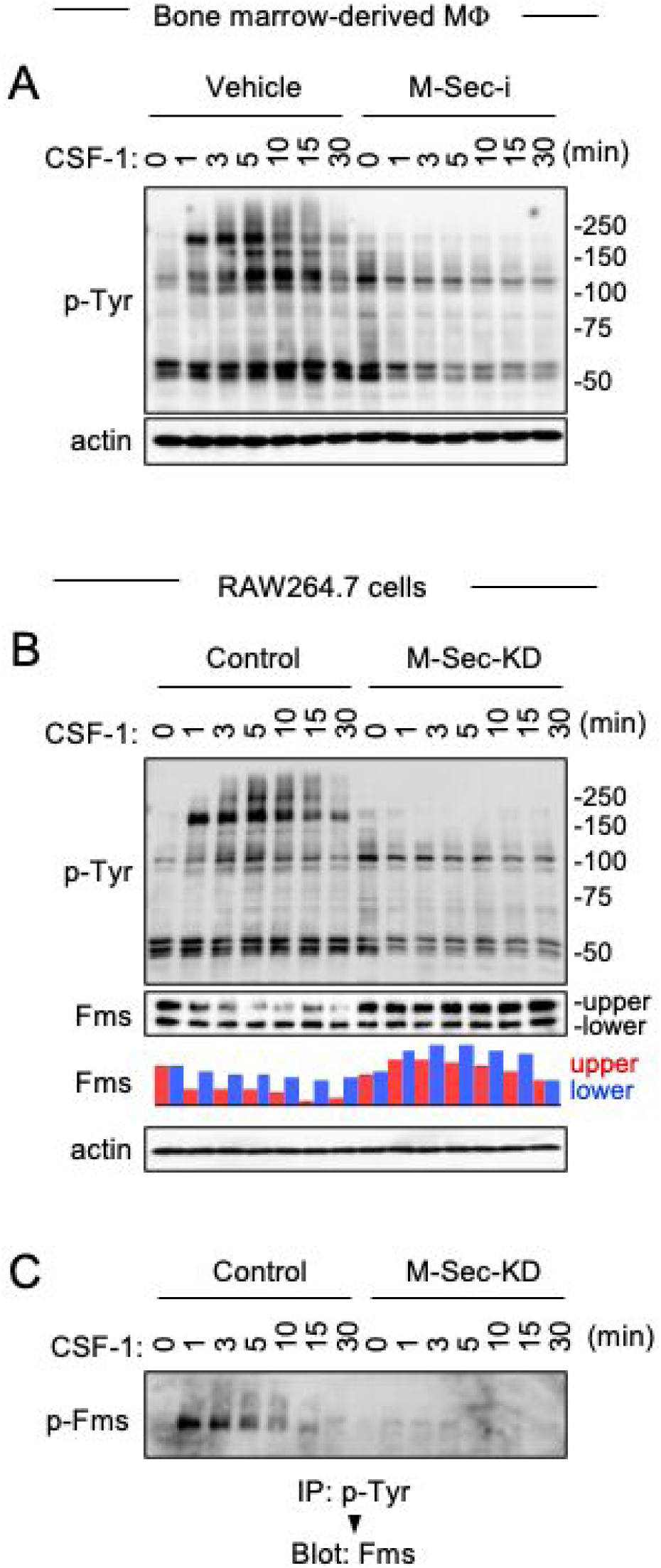
Effect of M-Sec inhibition or knockdown on CSF-1-induced Fms activation in macrophages. (A) Bone marrow-derived macrophages were pretreated with DMSO (Vehicle) or M-Sec inhibitor (M-Sec-i) for overnight, M-CSF starved for 6 hours, and left untreated or treated with CSF-1 for the indicted periods. The total cell lysates were subjected to western blotting using anti-phosphotyrosine (p-Tyr) antibody. β-actin blot is the loading control. (B) The control or M-Sec knockdown (M-Sec-KD) RAW264.7 cells were serum starved for 12 hours, and left untreated or treated with CSF-1 for the indicated periods. The total cell lysates were subjected to western blotting using anti-phosphotyrosine (p-Tyr) or anti-Fms antibody. The density of upper (red) or lower (blue) band of Fms is summarized (see text for details of the upper and lower Fms bands). The level shown is relative to that of untreated cells. β-actin blot is the loading control. (C) The control or M-Sec knockdown (M-Sec-KD) RAW264.7 cells were serum starved for 12 hours, and left untreated or treated with CSF-1 for the indicated periods. The anti-phosphotyrosine (p-Tyr) immunoprecipitates (IP) were analyzed by western blotting (Blot), using anti-Fms antibody to detect phosphorylated Fms (p-Fms).

There were two Fms bands in unstimulated cells, and the upper band decreased after CSF-1 stimulation in the control cells (Figure 5B, Fms blot and bar graph). This is because the upper band is the fully *N*-glycosylated cell surface form whereas the lower is the hypo-*N*-glycosylated immature intracellular form (32), and Fms activation associates with its endocytosis and degradation (33). Importantly, such Fms degradation was modest in the M-Sec knockdown cells (Figure 5B, Fms blot and bar graph). Consistent with this, the amount of tyrosine-phosphorylated Fms after CSF-1 stimulation, which reflects the activation and auto-phosphorylation of Fms, was obvious in the control cells, but weak in the M-Sec knockdown cells (Figure 5C).

### Exogenous M-Sec expression augments Fms aggregate formation and CSF-1-mediated Fms activation in 293 cells

To further confirm the effects of M-Sec on the CSF-1/Fms axis, we next performed the exogenous expression experiments, using 293 cells which are negative for endogenous expression of M-Sec and Fms. M-Sec expression in 293 cells augmented the formation of large Fms aggregates (Figure 6A) and increased the size of Fms aggregates (Figure 6B). Furthermore, the M-Sec expression augmented CSF-1-mediated activation of MAP kinases (Figure 6C) and phosphorylation of Fms at Tyr809 (Figure 6D), which is the major auto-phosphorylation site important for full activation of Fms (34). These results supported the idea that M-Sec was required for large Fms aggregate formation and important for the efficient CSF-1-induced activation of Fms.

**Figure 6.**
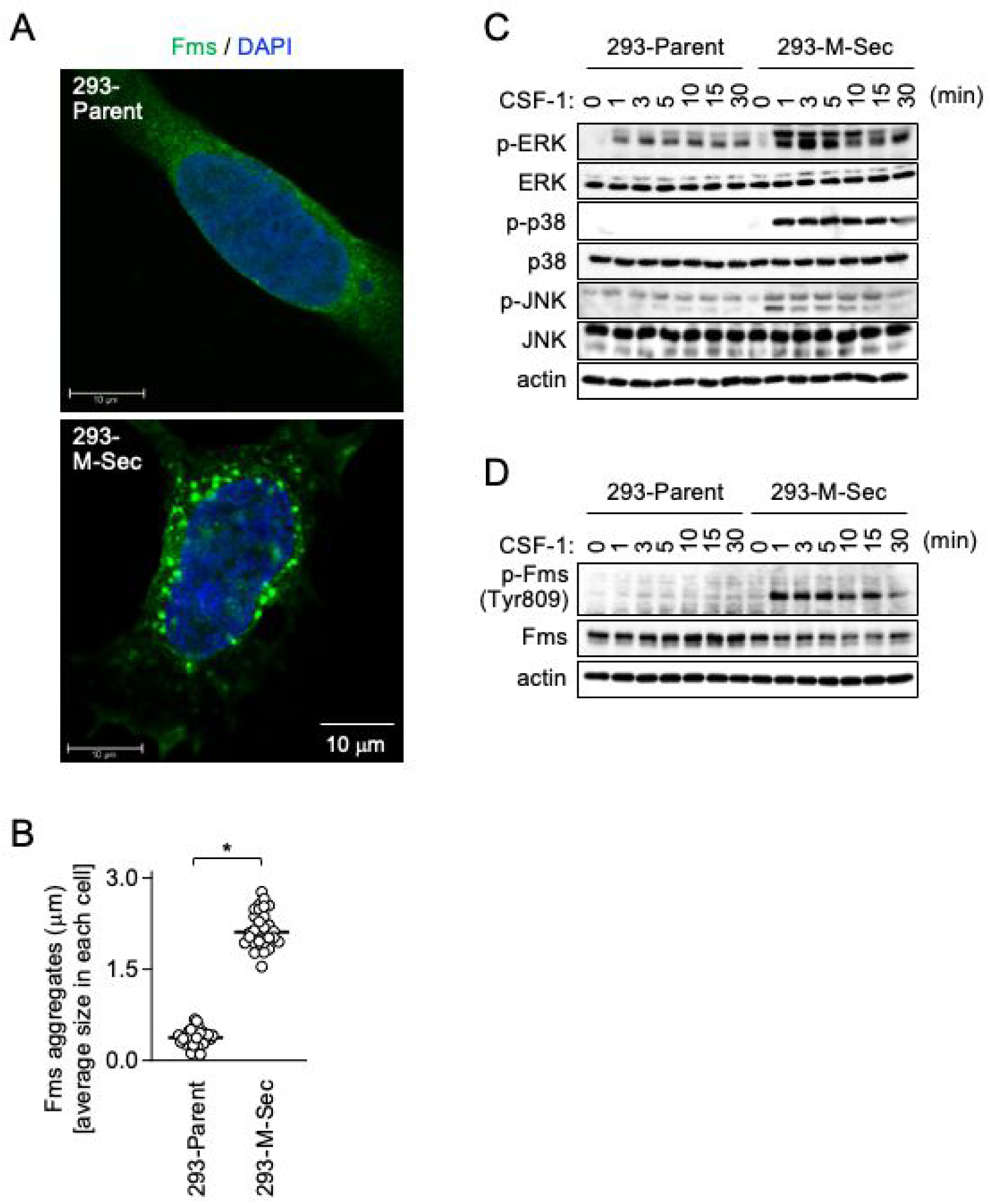
Effect of M-Sec expression on Fms aggregate formation and CSF-1-induced Fms activation in 293 cells. (A) The parent or M-Sec-expressing 293 cells were transected with the wild-type Fms plasmid, cultured for 2 days, co-stained with anti-Fms antibody (green) and DAPI (blue), and analyzed by immunofluorescence. Scale bar: 10 μm. (B) The parent or M-Sec-expressing 293 cells were analyzed as in **A**, and the average size of Fms aggregates in each cell is summarized (30 cells for each group). \**p* < 0.05. (C) The parent or M-Sec-expressing 293 cells were serum starved for 12 hours, and left untreated or treated with CSF-1 for the indicted periods. The total cell lysates were subjected to western blotting. Antibodies used were as follows: anti-phosphorylated ERK (p-ERK), anti-total ERK, anti-phosphorylated p38 (p-p38), anti-total p38, anti-phosphorylated JNK (p-JNK), and anti-total JNK. β-actin blot is the loading control. (D) The total cell lysates of the parent or M-Sec-expressing 293 cells were prepared as in **C**, and analyzed by western blotting using the antibody specific for tyrosine809-phosphorylated Fms (p-Fms). Total Fms was also analyzed. β-actin blot is the loading control.

### Fms mutant lacking motif that mediates binding to phosphatidylinositol 4,5-biphosphate fails to form large aggregates

We next attempted to clarify how M-Sec regulates Fms aggregate formation. M-Sec is known to bind phosphatidylinositol 4,5-biphosphate (PIP2), which is the most abundant phosphoinositide and present in the inner leaflet of the plasma membrane (35–37), via its N-terminal lysine-rich motifs (25). The similar lysine-rich motif is found in the intracellular juxtamembrane region of many receptor tyrosine kinases including Fms (38). EGF receptor, the best characterized receptor tyrosine kinase, is supposed to bind PIP2 through its lysine-rich motif (39). Thus, we next performed a series of experiments to test the hypothesis that PIP2 is required for M-Sec to augment Fms aggregate formation.

Fms has the lysine-rich putative PIP2-binding motif in its juxtamembrane region (^538^YKYKQKPK^545^, Figure 7A) (38). In M-Sec-expressing 293 cells, the 1-545 mutant, which lacked the most cytoplasmic domain but had the ^538^YKYKQKPK^545^ sequence, formed large aggregates (Figure 7A, upper), as the wild-type Fms did (see Figure 6A, lower). In contrast, the 1-537 mutant, which completely lacked the intracellular domain including the ^538^YKYKQKPK^545^ sequence, failed to form such large aggregates (Figure 7B, lower). This difference between these mutants was confirmed by the quantification of the size of Fms aggregates (Figure 7C).

**Figure 7.**
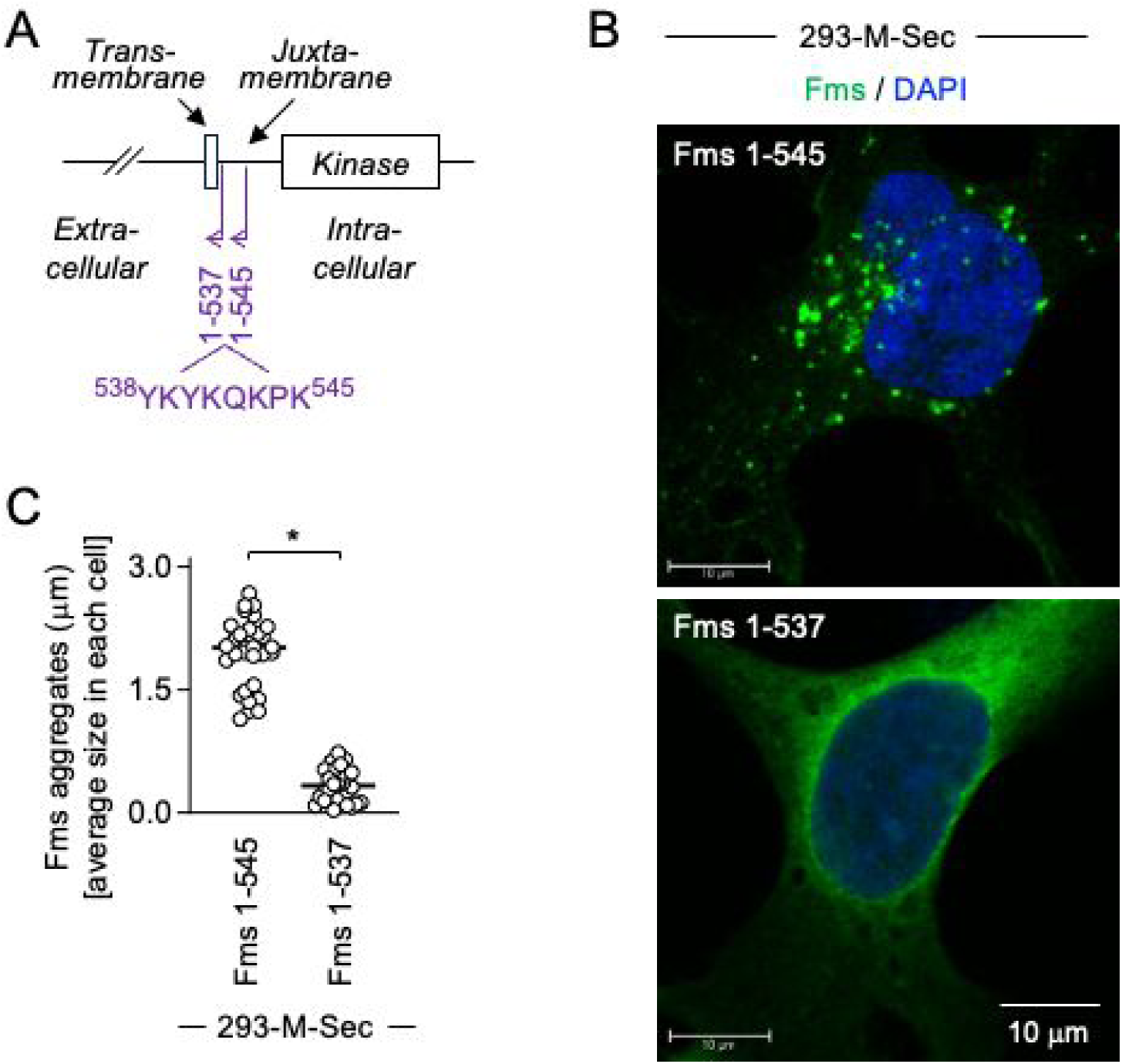
Aggregate formation of Fms mutants in M-Sec-expressing 293 cells. (A) The C-terminally deleted Fms mutants (1-537 and 1-545) are schematically shown. The amino acid sequence (^538^YKYKQKPK^545^) of the lysine-rich putative PIP2-binding motif is also shown. (B) The M-Sec-expressing 293 cells were transfected with the 1-545 or 1-537 mutant Fms plasmid, cultured for 2 days, co-stained with anti-Fms antibody (green) and DAPI (blue), and analyzed by immunofluorescence. Scale bar: 10 μm. (C) The M-Sec-expressing 293 cells were analyzed as in **B**, and the average size of Fms aggregates in each cell is summarized (30 cells for each group). \**p* < 0.05.

### M-Sec mutant lacking PIP2-binding motif fails to augment large Fms aggregate formation

M-Sec has two lysine-rich motifs at its N-terminus (see Figure 8A), and both the mutant lacking the first motif (ΔK1) and the mutant lacking the second motif (ΔK2) lost the binding to PIP2 (25). Thus, we established 293 cells expressing M-Sec ΔK1 or ΔK2 (Figure 8B). These cells expressed Fms, the level of which was comparable to that of 293 cells expressing the wild-type M-Sec (Figure 8C). However, the size of Fms aggregates in these cells was markedly smaller than that of cells expressing the wild-type M-Sec (Figure 8D).

**Figure 8.**
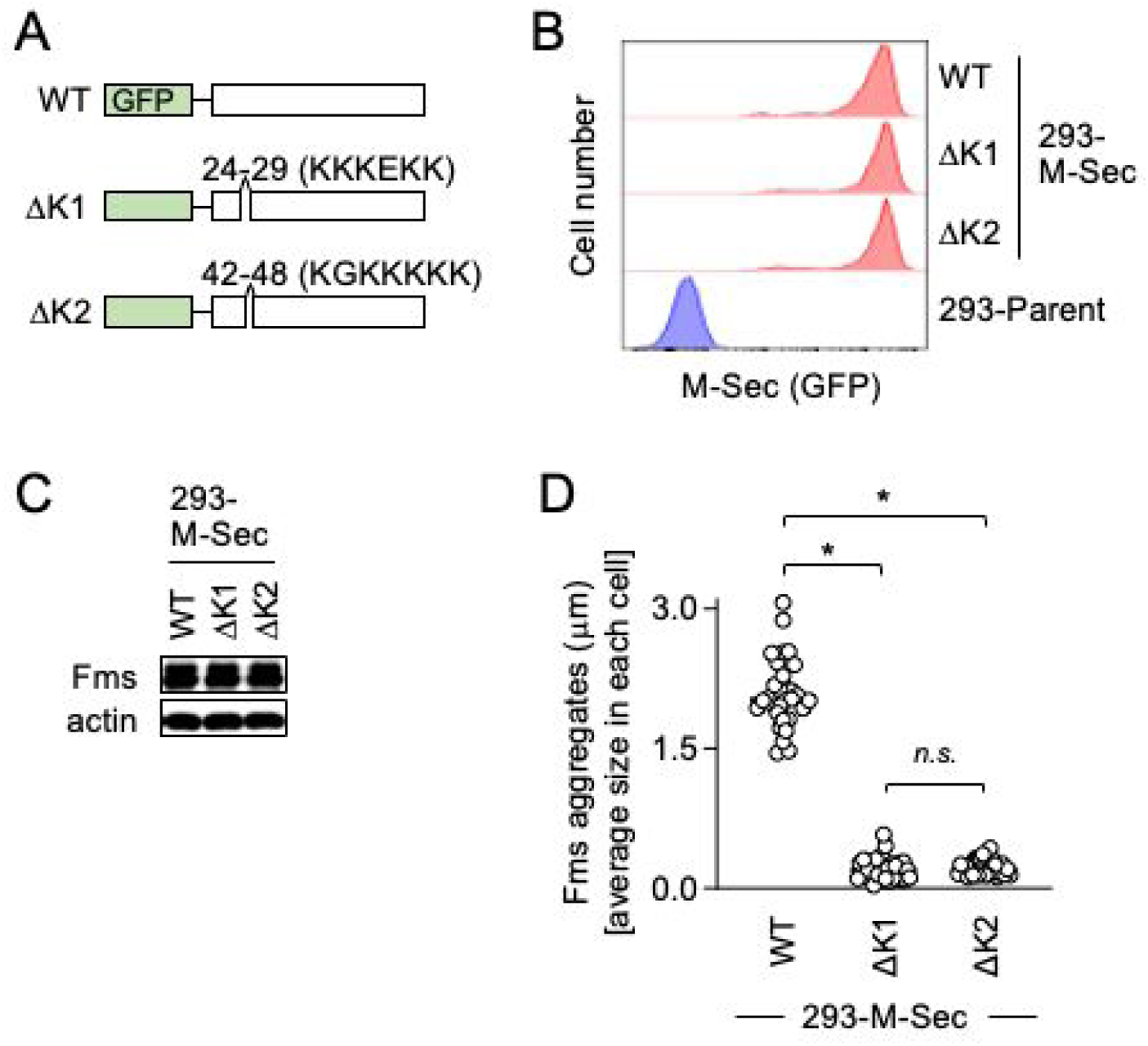
Fms aggregate formation in mutant M-Sec-expressing 293 cells. (A) The GFP-fused wild-type (WT) M-Sec and its mutants (ΔK1 and Δκ2) used are schematically shown. The ΔK1 and Δκ2 lack the first (^24^KKKEKK^29^) and second lysine-rich PIP2-binding motif (^42^KGKKKKK^48^), respectively. (B) The 293 cells were engineered to stably express M-Sec WT, ΔK1 or Δκ2. The expression of these GFP-fused M-Sec proteins was confirmed by flow cytometry. The parent 293 cells were also analyzed as a reference. (C) The 293 cells expressing M-Sec WT, ΔK1 or Δκ2 were transfected with Fms plasmid, cultured for 2 days, and analyzed for the expression level of Fms by western blotting. β-actin blot is the loading control. (D) The 293 cells expressing M-Sec WT, ΔK1 or Δκ2 were transfected with Fms plasmid, cultured for 2 days, stained with anti-Fms antibody, and analyzed by immunofluorescence. Then, the average size of Fms aggregates in each cell is summarized (30 cells for each group). *n.s.*, not significant. \**p* < 0.05.

### Depletion of cellular PIP2 reduces M-Sec-mediated large Fms aggregate formation

To further confirm the involvement of PIP2 in M-Sec-mediated large Fms aggregate formation, we performed the experiment using the plasmid expressing inositol polyphosphate 5-phosphatase type IV (5ptase), which reduces cellular level of PIP2 (Ref. 40, 41). When expressed in 293 cells expressing M-Sec, 5ptase did not affect Fms expression (Figure 9A), but reduced the formation of large Fms aggregates (Figure 9B) and the size of Fms aggregates (Figure 9C). These results, together with the results of experiments using Fms mutants (see Figure 7) and M-Sec mutants (see Figure 8), strongly suggest that PIP2 is required for M-Sec to augment Fms aggregate formation.

**Figure 9.**
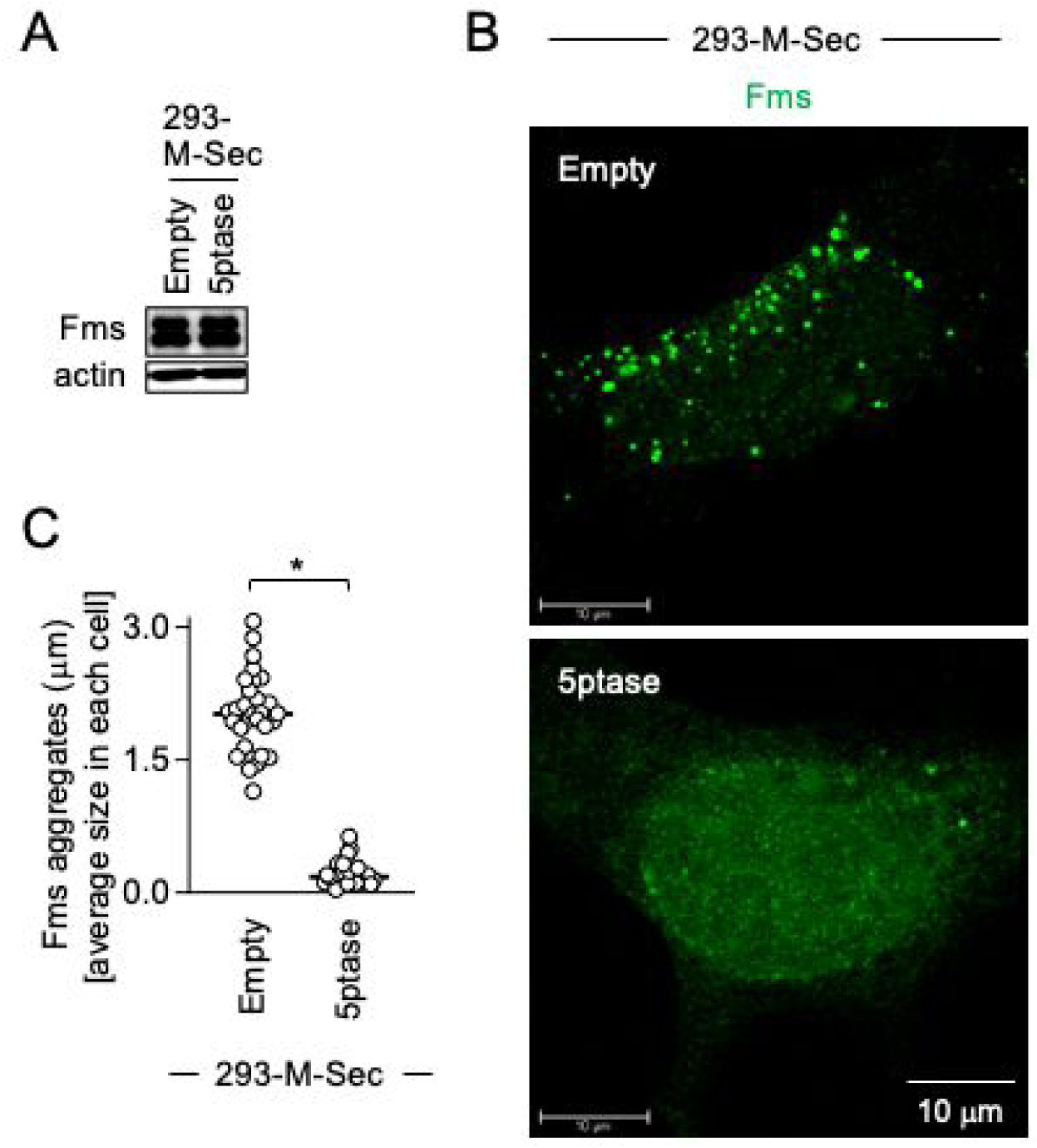
Effect of the expression of inositol polyphosphate 5-phosphatase type IV on Fms aggregate formation. (A) The M-Sec-expressing 293 cells were co-transfected with Fms plasmid and either the empty plasmid (Empty) or inositol polyphosphate 5-phosphatase type IV plasmid (5ptase). Then, the cells were analyzed for the expression level of Fms by western blotting. β-actin blot is the loading control. (B) The M-Sec-expressing 293 cells were co-transfected with Fms plasmid and either the empty (Empty) or 5ptase plasmid (5ptase). Then, the cells were cultured for 2 days, stained with anti-Fms antibody (green), and analyzed by immunofluorescence. Scale bar: 10 μm. (C) The M-Sec-expressing 293 cells were analyzed as in **B**, and the average size of Fms aggregates in each cell is summarized (30 cells for each group). \**p* < 0.05.

### M-Sec alters cellular distribution of PIP2

Finally, we performed the experiment using the plasmid expressing GFP-fused pleckstrin homology domain of phospholipase Cδ (GFP-PLCδ-PH), which is widely used as a probe to visualize cellular PIP2 distribution in living cells (41–43). Interestingly, parental 293 cells and M-Sec-expressing 293 cells showed a marked difference in the distribution of GFP-PLCδ-PH (Figure 10A): the large GFP-PLCδ-PH-positive structures were observed in M-Sec-expressing 293 cells (lower panels). There was no difference in the percentage of GFP-PLCδ-PH-positive cells or expression level of GFP-PLCδ-PH between parental 293 cells and M-Sec-expressing 293 cells (Figure 10B). These results suggest that M-Sec alters cellular distribution of PIP2, which leads to the formation of large Fms aggregates.

**Figure 10.**
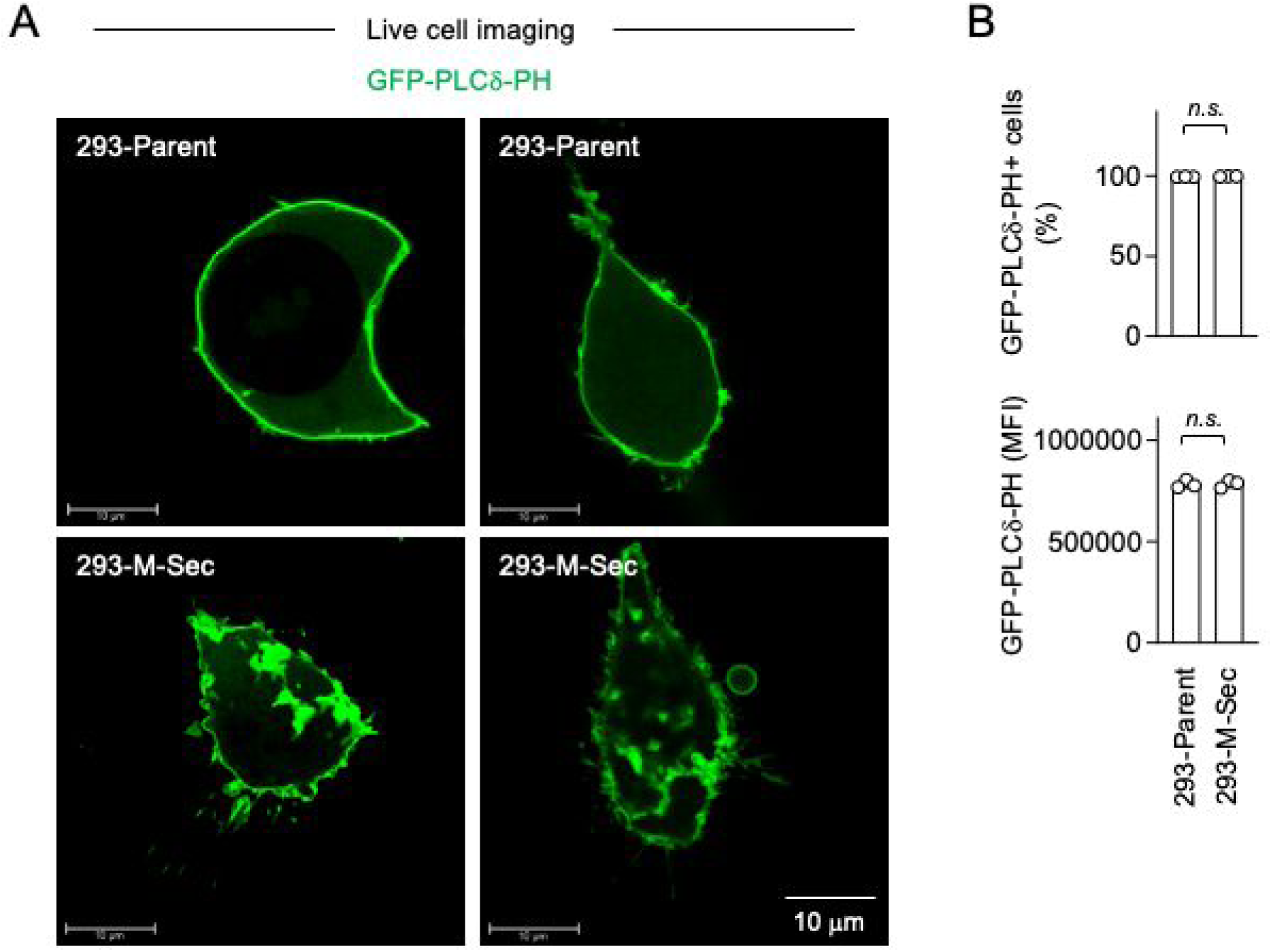
Effect of M-Sec on cellular distribution of PIP2. (**A**, **B**) The parent or M-Sec-expressing 293 cells were transfected with GFP-PLCδ-PH plasmid, and cultured for 2 days. In **A**, the cells were analyzed for GFP expression by live cell imaging to visualize cellular distribution of PIP2. Scale bar: 10 μm. In B, the cells were analyzed for GFP expression (to detect GFP-PLCδ-PH) by flow cytometry (n=3). The percentage of GFP-positive cells and mean fluorescence intensity (MFI) are summarized in the left and right panels, respectively. *n.s.*, not significant.

## DISCUSSION

In this study, we identified an unreported function of M-Sec in macrophages. We revealed that M-Sec is required for the efficient functional and signaling response of macrophages to CSF-1, but not to CSF-2. In fact, M-Sec is required for the efficient CSF-1-induced activation of its receptor Fms. We also revealed that M-Sec is required for the formation of large Fms aggregates, which have been supposed to be beneficial for CSF-1 to dimerize and activate Fms (21). The cellular phosphoinositide PIP2 is required for the formation of the large Fms aggregates by M-Sec. Collectively, it is likely that M-Sec regulates cellular localization of Fms via PIP2 and augments CSF-1-induced activation of Fms. M-Sec appears a unique cellular regulator of Fms activation.

In the absence of CSF-1 or IL-34, Fms is present as the inactive monomer due to cis-autoinhibition (16, 17). The binding of CSF-1 or IL-34, both of which are homodimers, induces the dimerization of Fms and releases the cis-autoinhibition (16–18). Thus, the dimerization is critical step for the activation of Fms, and the abundance or distribution of Fms at the plasma membrane is the rate-limiting step in its activation. Because of this, the large Fms aggregates in which the monomers are close to each other are supposed to be beneficial for CSF-1 or IL-34 to dimerize Fms (21). Our findings suggest that M-Sec brings the Fms monomers close to each other via PIP2 and therefore facilitates the efficient dimerization and activation of Fms in response to CSF-1 or IL-34.

The present study at least in part answers the two long-standing questions, namely, the mechanisms by which Fms forms large aggregates in macrophages and the significance of the large Fms aggregates for CSF-1-induced Fms activation. Apart from its physiologically high expression in macrophages, M-Sec is aberrantly expressed in nasopharyngeal carcinoma cells (44). Interestingly, Liu and Wu recently reported that M-Sec knockdown in nasopharyngeal carcinoma cells reduced the auto-phosphorylation of the receptor tyrosine kinase EGF receptor in response to its ligand EGF (45; meeting abstract). Although the details including underlying mechanisms are unclear, it is possible that the reduced cellular response to EGF by M-Sec knockdown is due to altered cellular localization of EGF receptor, as exemplified by Fms. In fact, EGF receptor is supposed to bind PIP2 through its lysine-rich motif (39). Thus, M-Sec may regulate not only Fms but also other receptor tyrosine kinases including EGF receptor.

HTLV-1 is the causative virus of adult T-cell leukemia. We previously reported that HTLV-1 induces the aberrant expression of M-Sec in CD4^+^ T cells and that M-Sec inhibition or knockdown alters the cellular localization of Gag, the structural protein of HTLV-1 (28). Since Gag has a lysine- and arginine-rich polybasic motif at its N-terminus (46), it is possible that the altered Gag localization by the M-Sec inhibition/knockdown is related to the binding of Gag to PIP2 through the motif.

Our results suggest that M-Sec alters cellular distribution of PIP2, which leads to the formation of large Fms aggregates. However, how M-Sec alters cellular distribution of PIP2 is unclear. M-Sec is thought to initiate the formation of tunneling nanotubes through RalA, a small GTPase, and the exocyst complex as the downstream effector of RalA (24, 25), although the precise role of M-Sec, RalA and the exocyst complex is not fully understood. The exocyst complex is evolutionary conserved and known to tether secretory vesicles to the plasma membrane (47, 48). Among eight components of the exocyst complex, Sec3 (also known as EXOC1) and Exo70 (also known as EXOC7) bind PIP2 (49, 50). Thus, these components may help to alter cellular PIP2 distribution.

A number of cellular proteins have been proposed to bind PIP2 and induce the clustering of PIP2 (36). However, the binding and clustering may not be enough to explain the altered cellular PIP2 distribution induced by M-Sec because the PIP2 signals in the presence of M-Sec were very large (see Figure 10). The formation of tunneling nanotubes, the hallmark function of M-Sec, may help to induce such large PIP2 accumulation.

Jin at al. recently reported that M-Sec knockdown in macrophages reduced the activation of MAP kinases in response to H_2_O_2_ (51). However, the underlying mechanisms are unknown. Interestingly, H_2_O_2_ activates a Ca^2+^-permeable channel TRPM2 (formerly known as LTRPC2) (52), and PIP2 was proposed to regulate the activation of TRPM2 (53). Thus, the PIP2 may be involved in the reduced activation of MAP kinases in the M-Sec knockdown macrophages.

In this study, we used CSF-1 as the ligand of Fms and have demonstrated that M-Sec regulates the CSF-1/Fms axis. Because IL-34 is the alternative ligand of Fms, M-Sec may also regulate the IL-34/Fms axis. However, more studies are necessary to clarify whether the formation of large Fms aggregates via PIP2 is the sole mechanism by which M-Sec augments the functional and signaling response of macrophages to CSF-1, because PIP2 is known to support a broad spectrum of cellular functions and the best characterized regulator of actin cytoskeleton among all phosphoinositides (35–37). Despite the unresolved questions, M-Sec will be helpful to understand the mechanism of activation of Fms and other receptor tyrosine kinases including EGF receptor.

## MATERIALS AND METHODS

### Bone marrow-derived self-renewing macrophages

Mouse bone marrow-derived macrophages that stably proliferated in the presence of CSF-1 were established previously (31). They were maintained with RPMI1640 medium/10% FCS containing recombinant human (rh) CSF-1.

### RAW264.7 cells

The control and M-Sec knockdown mouse macrophage-like RAW264.7 cells were established previously (25). They were maintained with RPMI1640 medium/10% FCS. The control or M-Sec knockdown cells were further engineered to express human M-Sec. The human M-Sec cDNA (NCBI GenBank M92357.1) was synthesized at Eurofins Japan and cloned into the lentiviral vector pLVSIN-EF1α-Hyg (TaKaRa-Bio). The lentivirus was prepared and added to the control or M-Sec knockdown cells, and cells expressing human M-Sec were selected under the cultures containing 750 μg/ml hygromycin B.

### CSF-1 and CSF-2

rhCSF-1 was a kind gift from Morinaga Milk Industry (Kanagawa, Japan), and used at a final concentration of 100 ng/ml. The recombinant mouse (rm) CSF-2 (Miltenyi Biotec) was used at a final concentration of 10 ng/ml.

### M-Sec inhibitor

The M-Sec inhibitor was identified by an affinity-based chemical array screening (26). In brief, test compounds were arrayed onto the photoaffinity linker-coated slides, which were incubated with the lysates of 293 cells expressing either the control DsRed or DsRed-M-Sec fusion protein. The fluorescent signals were quantified, and the compound NPD3064 to which DsRed-M-Sec, but not DsRed, bound was identified. The M-Sec inhibitor was synthesized at Pharmeks (Moscow, Russia), dissolved in DMSO, and added to cultures at a final concentration of 10 μM (0.1% v/v). The same volume of DMSO was added as a vehicle control.

### 293 cells stably expressing M-Sec

The 293A cells (Invitrogen) were maintained with DMEM medium/10% FCS, and engineered to express human M-Sec using the lentiviral system. The lentivirus was added to 293 cells, and cells expressing human M-Sec were selected under the cultures containing 200 μg/ml hygromycin B. The 293A cells were also engineered to express GFP-fused mouse M-Sec. They were transfected with the plasmid encoding the wild-type or mutant (ΔK1 or ΔK2) (25), using Lipofectamine 3000 reagent (Invitrogen). The cells expressing GFP-fused M-Sec were selected under the cultures containing 1,200 μg/ml G418, and enriched by cell sorting using FACSAria II (BD Biosciences).

### Transfection using 293 cells

The 293 cells were seeded onto 12-wll plates, cultured, and transfected with various plasmids using 3 μl Lipofectamine 3000 reagent and 2 μl P3000 reagent (both from Invitrogen). The total amount (1 μg) of the plasmid was normalized using appropriate control (empty) vectors. After 6 hours of transfection, the culture medium was replaced with fresh medium. The cells were further cultured for 1 or 2 days and subjected to immunofluorescence, live cell imaging, flow cytometry, or western blotting.

### Plasmids

The C-terminally deleted Fms cDNAs (1-545 and 1-537) were prepared using PCR. The PCR products were cloned into pCR2.1 vector (Invitrogen) and the nucleotide sequences were verified using a BigDye Terminator v3.1 Cycle Sequencing Kit and ABI PRISM 3100 Genetic Analyzer (both from Applied Biosystems). The insert cDNAs were subcloned into pcDNA3.1 expression vector (Invitrogen). The plasmid of inositol polyphosphate 5-phosphatase type IV was prepared as described previously (41). The GFP-PLCδ-PH plasmid was obtained through Addgene (#21179).

### Immunofluorescence and live cell imaging

Immunofluorescence was performed as described previously (28). In brief, cells were fixed in 4% paraformaldehyde, permeabilized with 0.1% Triton X-100, and stained with anti-mouse Fms (AFS98; BioLegend) or anti-human Fms antibodies (9-4D2-1E; BioLegend) followed by AlexaFluor488-labeled anti-rat IgG or AlexaFluor633-labeled anti-rat IgG (both from Molecular Probes). Cells were also stained with DAPI (Molecular Probes) to visualize nuclei. Signals were visualized using confocal laser-scanning microscope FV1200 (Olympus) or TCS SP8 (Leica). Image processing was performed using FV viewer ver. 4.2 software (Olympus) or LAS X software ver.1.4.5 (Leica). The signal of Fms aggregates was quantified using ImageJ 1.52n software (NIH). To visualize PIP2 in living cells, the signal of GFP-PLCδ-PH in unfixed 293 cells was analyzed using TCS SP8.

### Flow cytometry

The cell surface expression of Fms was analyzed by flow cytometry. In brief, cells were detached using enzyme-free cell dissociation buffer (Millipore), stained with fluorescent dye-labeled antibodies, and analyzed on FACSCanto II (BD Biosciences) or CytoFLEX (Beckman Coulter) using FlowJo software (FlowJo LLC). The antibodies used were as follows: allophycocyanin (APC)-labeled anti-mouse Fms (AFS98) and APC-labeled anti-human Fms (9-4D2-1E4) (both from BioLegend).

### Western blotting

Western blotting was performed as described previously (27). In brief, cells were lysed with Nonidet P-40 lysis buffer containing protease- and phosphatase inhibitors. In a selected experiment, the total cell lysates were incubated with anti-phosphotyrosine antibody (PY99) followed by Protein A/G PLUS Agarose (both from Santa Cruz Biotechnology). The total cell lysates or the anti-phosphotyrosine immunoprecipitates were subjected to western blotting. The antibodies used were as follows: anti-phospho ERK (E-4), anti-total ERK (C-9) (both from Santa Cruz Biotechnology), anti-phospho p38 (3D7; Cell Signaling Technology), anti-total p38 (A-12; Santa Cruz Biotechnology), anti-phospho JNK (81E11), anti-total JNK (#4672) (both from Cell Signaling Technology), anti-phosphotyrosine (PY99; Santa Cruz Biotechnology), anti-Fms (#3152; Cell Signaling Technology) (to detect mouse Fms), anti-Fms (E7S2S; Cell Signaling Technology) (to detect human Fms), anti-phospho Fms (Tyr809; Cell Signaling Technology), and anti-actin (EPR16769; Abcam). Detection was performed using HRP-labeled secondary antibodies (GE Healthcare), western blot ultra-sensitive HRP substrate (TaKaRa-Bio), and ImageQuant LAS4000 image analyzer (GE Healthcare). The density of the upper or lower band of Fms in RAW264.7 cells was quantified using the ImageJ software after normalization to the density of the actin band.

### Nitric oxide production and iNOS mRNA expression

The concentration of nitric oxide in the culture supernatants of RAW264.7 cells was quantified using Griess reagent (Dojindo, Kumamoto, Japan). The expression of iNOS mRNA was analyzed by real-time RT-PCR (31). In brief, RNA was isolated using ISOGEN II reagent (Nippon Gene, Tokyo, Japan), cDNA was prepared using M-MLV RT (Invitrogen), and real-time PCR was performed using SYBR Premix Ex Taq II (TaKaRa-Bio) and LightCycler (Roche). β-actin mRNA was quantified as an internal control. The primer pair for iNOS mRNA was as follows: 5’ ACCTTGTTCAGCTACGCCTT 3’ and 5’CATTCCCAAATGTGCTTGTC 3’.

### Phagocytosis assays

Phagocytic activity was quantified as described previously (54). In brief, RAW264.7 cells were incubated with fluorescent microspheres (fluoresbrite carboxylate microspheres with a 0.7 μm diameter, Polysciences) for 2 hours, and analyzed by flow cytometry as described above.

### Statistical analysis

Differences between two groups were analyzed by unpaired Student’s *t*-test. Differences between multiple groups were analyzed by a one-way or two-way ANOVA. All statistical analyses were conducted using PRISM 10 (GraphPad).

## Acknowledgments

We thank K. Nasu and I. Suzu for their technical and secretarial assistance, respectively.

## Author contributions

Conceptualization: SS. Investigation: RAA, MH, NT, YME. Methodology: HM. Resources: SK, KH, HO, KM, AO, RAA, MH. Writing-original draft: SS, RAA. Writing-review & editing: SK, KH, HO, KM, AO.

## Data availability

All data described in this manuscript are present in the manuscript.

